# Reduced enteric BDNF-TrkB signaling drives stress-dependent glucocorticoid-mediated GI dysmotility

**DOI:** 10.1101/2024.12.13.628260

**Authors:** Jared Slosberg, Srinivas N. Puttapaka, Philippa Seika, Su Min Hong, Alpana Singh, Ainsleigh Scott, Subhash Kulkarni

**Author notes:** Address all correspondence to: Subhash Kulkarni, Ph.D. Joint first co-authors (Jared Slosberg, Srinivas N. Puttapaka).

## Abstract

Stress is a key contributor to gastrointestinal (GI) dysmotility, particularly in patients with disorders of gut-brain interactions (DGBI). Since GI motility is governed by the enteric nervous system (ENS), stress may act by altering ENS function. While stress activates glucocorticoid signaling via the hypothalamic-pituitary-adrenal axis, the impact of stress-mediated glucocorticoid signaling on ENS biology remains poorly understood. In the central nervous system, glucocorticoids reduce specific isoforms of brain-derived neurotrophic factor (BDNF), impairing signaling through its receptor, TrkB, and contributing to behavioral dysfunction. However, the identity of ENS-specific *Bdnf* isoforms, their glucocorticoid sensitivity, and the effect of enhanced TrkB signaling on GI motility in stressed animals has not been characterized. Here, using male and female mice, we show that >85% of post-natal ENS *Bdnf* transcripts are glucocorticoid-responsive isoforms. We also demonstrate that both BDNF and its receptor TrkB (*Ntrk2*) are expressed by enteric neurons. In male mice, stress and administration of dexamethasone—a synthetic glucocorticoid receptor (GR) agonist—cause GI dysmotility, which we demonstrate is associated with significantly reduced *Bdnf* transcripts in the longitudinal muscle – myenteric plexus (LM-MP) tissue *in vivo*. Dexamethasone exposure also represses *Bdnf* transcript and mature protein levels in LM-MP tissue *in vitro*. Notably, treatment with HIOC, a selective TrkB agonist, rescues GI transit defects in dexamethasone-treated animals. These findings identify BDNF-TrkB signaling as a key modulator of stress-induced ENS dysfunction and highlight TrkB as a promising therapeutic target for GI dysmotility in DGBI.

**Significance statement:** How stress causes gastrointestinal (GI) dysmotility is not well understood. GI motility is regulated by the enteric nervous system (ENS), which is responsive to brain-derived neurotrophic factor (BDNF), which signals through its receptor tropomyosin related kinase B (TrkB). By altering glucocorticoid signaling, stress modulates brain’s BDNF levels to cause behavioral changes. However, if this pathway is similarly responsible for stress’s effects on GI dysmotility is not well understood. Here, by identifying the nature of ENS-specific *Bdnf* isoforms, studying their response to stress and glucocorticoid signaling, and testing the effect of a TrkB agonist to improve gut motility in a model of glucocorticoid-driven dysmotility, we implicate altered BDNF-TrkB signaling as an important mechanism driving stress-associated dysmotility.

## Introduction

Stress-associated gastrointestinal (GI) dysmotility, a prevalent issue in irritable bowel syndrome (IBS), afflicts large numbers of patients (Leigh *et al*, 2023; Qin *et al*, 2014; Sperber *et al*, 2021). GI motility is regulated by the enteric nervous system (ENS) (Wood, 2016), suggesting that stress-associated GI dysmotility can occur due to ENS dysfunction. Healthy ENS function depends on optimal neurotrophin signaling, where neurotrophins secreted by various gut cell types interact with their receptors on ENS neurons (Chalazonitis *et al*, 2020; Liu, 2018; Sharkey & Mawe, 2023). Brain-derived neurotrophic factor (BDNF) and its receptor tropomyosin kinase B (TrkB, coded by *Ntrk2*) comprise of one such system. In the central nervous system (CNS), stress reduces *Bdnf* expression, impairing BDNF – TrkB signaling to alter CNS neuronal function and organisms’ behavior (Martinowich et al., 2007; Zheng et al., 2024; Begni et al., 2016). However, it remains unclear whether stress-associated GI dysmotility involves reduced *Bdnf* expression, and if so, by what mechanisms stress alters enteric *Bdnf* expression

Rodent studies have demonstrated that BDNF regulates GI motility, as its presence hastens GI transit of luminal contents (Boesmans *et al*, 2008; Chen *et al*, 2014; Chen *et al*, 2012; Singh *et al*, 2020). Mirroring this, decreased BDNF was detected in colonic tissues from patients with slow-transit constipation (Chen *et al*., 2014), and exogenous supplementation of BDNF improved GI motility in constipated patients (Coulie *et al*, 2000). These data suggest that mechanisms regulating *Bdnf* expression are relevant for pathobiology of GI dysmotility. Further, altered *Bdnf* transcript levels in distressed IBS patients (Konturek *et al*, 2020; Zhang *et al*, 2021) suggest that *Bdnf* dysregulation may extend to stress-induced GI dysmotility. However, how stress affects *Bdnf* expression in GI motility-regulating cells, and whether normalizing BDNF-TrkB signaling in stressed animals rescues GI motility is not well understood.

The *Bdnf* gene consists of 8 non-coding (e1-e8) and 1 coding (e9) exons (Aid *et al*, 2007). Each non-coding exon possesses a unique promoter, allowing for differential regulation of various *Bdnf* isoforms (Aid et al, 2007). Variation in promoter usage underlies context-dependent regulation of *Bdnf* expression (Hong *et al*, 2008; Pruunsild *et al*, 2011). One of these contexts, stress, represses the expression of two specific isoforms – *Bdnf-IV* and *Bdnf-VI* – through hypothalamic-pituitary-adrenal axis (HPA)-dependent modulation of glucocorticoid (GCs) levels (Hill & Martinowich, 2016; Linz *et al*, 2019; Murakami *et al*, 2005; Tsankova *et al*, 2006). GCs activate glucocorticoid receptor (GR), which then bind to glucocorticoid responsive elements (GREs) in the promoters of exons (e4 and e6) of *Bdnf-IV* and *Bdnf-VI* isoforms to downregulate their expression (Chen *et al*, 2017). While stress and GC supplementation reduce gut motility in adult mice (Schneider *et al*, 2023), it remains unknown if repression of specific *Bdnf* isoforms in the ENS underlies stress-related GI dysmotility. Interrogating this biology has been limited by the unclear identity of BDNF- and TrkB-expressing cells in the gut wall, alongside a lack of profiling of *Bdnf* isoform expression patterns and their susceptibility to stress and GC stimulation.

To address these gaps, here, we used immunofluorescence, bulk transcriptomics, and quantitative PCR to study the expression of BDNF and TrkB in the longitudinal muscle containing myenteric plexus (LM-MP) tissue, which contains the bulk of the motility-regulating ENS neurons. Further, we used restraint stress and dexamethasone, a synthetic GR agonist, to assess how stress and GR stimulation influence *Bdnf* expression to drive GI dysmotility and whether stimulating TrkB signaling rescues GI motility in a mouse model of GC-induced GI dysmotility. Our results show BDNF and TrkB are expressed by adult enteric neurons, and that stress-responsive isoforms represent the majority of *Bdnf* isoforms in the post-natal myenteric plexus tissue. We find that stress and dexamethasone exposure reduce expression of stress-responsive *Bdnf* isoforms and cause GI dysmotility - which is reversed by using HIOC, a selective agonist of BDNF-receptor TrkB.

## Methods

### Human tissue procurement and immunohistochemistry

Paraffin sections of the adult small intestinal full-thickness gut from deidentified human tissues were obtained from the Department of Pathology at Beth Israel Deaconess Medical Center, and the Institutional Review Board approved using the deidentified human tissues. The tissues were obtained from adult patients without any known motility disorder, and the deidentified nature of specimens preclude us from providing further identifying information. Antigen retrieval was performed on 5 µm thick sections at 97°C for 40 minutes using citrate buffer at pH 6.0, followed by immunostaining with anti-BDNF antibody (abcam, Cat. No. ab108319, dilution 1:500). The tissue sections were then counterstained with DAB chromogen staining, mounted with Cytoseal, and imaged under 40X brightfield. Microscopy was performed using the Nikon Eclipse Ni-U Upright microscope with 40x Nikon Plan Fluor objective (NA: 0.75). Images were acquired using the Nikon DS-Fi3 camera and NIS-Elements BR software (Version 5.20.01). During image acquisition, exposure time was adjusted using the Color DS-Fi3 settings to obtain a clear definition of tissue structure. Images were acquired in the TIFF format.

### Animals

We used male and female wildtype C57BL/6J mice from the National Institute on Aging colony (at the ages of 1, 6, and 17 months) for the bulk RNA sequencing experiment. For all other experiments – except for the immunostaining to quantify proportions of BDNF^+^ or TrkB^+^ neurons, we used adult male wildtype C57BL/6J mice (between the ages of 12 and 20 weeks) from the National Institute on Aging colony. For experiments to qualitatively and quantitatively assess nature of BDNF-expressing and TrkB-expressing neurons, we used 2 – 4 months old male TrkB-GFP mice (kind gift from Dr. David Ginty, Harvard), which have been previously used to observe TrkB-expressing neurons in the myenteric plexus (Nestor-Kalinoski *et al*, 2022). For testing the effect of loss of TrkB (*Ntrk2*) in the ENS on GI function, *Wnt1*-cre mice, which have been previously used by us to label all neural crest-derived neurons and glial cells in the adult ENS, were crossed with TrkB^fl/WT^ mice (Jax Strain #022363, kind gift from Dr. David Ginty, Harvard University) to generate the *Wnt1*-cre:TrkB^fl/fl^ (cause a conditional loss of TrkB from neural crest-derived neurons) and *Wnt1*-cre:TrkB^WT/WT^ (control) mice. Mice were housed in a controlled environment under a 12 h light/dark cycle at 22 ± 2℃, 55 ± 5% humidity with access to food and water ad libitum. All animal procedures were carried out strictly under protocols approved by the Animal Care and Use Committee of Johns Hopkins University and of Beth Israel Deaconess Medical Center (BIDMC) in accordance with the guidelines provided by the National Institutes of Health.

### Bulk RNA sequencing (bulk RNA-seq) experiments

#### Tissue Isolation for bulk RNA-seq

Mice of three ages (1-month, 6-month, and 17-month) and both sexes (n=3 per unique condition) were anesthetized with isoflurane and sacrificed by cervical dislocation. For small intestinal tissue isolation, a laparotomy was performed, and the entire small intestine was removed and flushed with ice-cold nuclease-free 1X PBS. The ileum, defined as the distal third of the small intestine, was cut into 3 cm sections for peeling. Next, tissues were placed over a sterile plastic rod and a superficial longitudinal incision was made along the serosal surface and the longitudinal muscle containing myenteric plexus (LM-MP) was peeled off from the underlying tissue using a wet sterile cotton swab. Each gut muscle strip was immediately flash frozen in liquid nitrogen before storage at −80°C. The average time from sacrifice to flash freeze was 10 minutes. Animals were processed in three batches, balanced across age/sex composition and order of processing.

#### Library Prep & Sequencing

RNA isolation, library preparation, and sequencing were completed at the JHU Single Cell and Transcriptomic Core. Samples with RIN >= 7.0 were used for library preparation, with all samples being high-quality except one 17-month female (resulting n=2, average RIN of 17 samples that passed this threshold > 9.0). Frozen tissue samples each in 1.5mL tubes were submerged in 375 ul of RLT buffer with β-mercaptoethanol. Tissue and buffer were transferred to an ice-cold 2 mL Qiagen sample tube with a 5mm stainless steel bead and lysed in a TissueLyzer at full speed for 3 minutes. RNA isolation was completed with the Qiagen RNeasy Mini, Animal Tissue and Cells with DNA Digest protocol. cDNA libraries were prepared with the TruSeq Stranded mRNA Library Prep with unique dual indexes. Paired-end (2x-50bp) sequencing was completed on a NovaSeq 6000 system.

#### Preprocessing

Isoform-level transcript abundances were estimated by pseudoalignment via Kallisto ‘quant’ (version 0.48.0) to the GENCODE vM27 reference genome assembly. Default parameters were used, except the number of bootstraps was set to 100 to estimate uncertainty (“-b 100”). The median fragment length, as estimated by Kallisto, was 168.7 base pairs. Across all sequenced samples, the median number of reads per sample was 46,756,241 (range: 34.6 M to 58.2 M) and 94.2% of reads were successfully pseudo-aligned to the reference.

#### Quality control & analysis

For quality control analyses, isoforms were collapsed to gene-level and abundances were normalized by the “scaledTPM” option within tximport, and further analyzed with DESeq2 (v1.30.1, R v4.0.2). Genes were removed from the analysis if they had fewer than 10 counts across all samples or were not detected in at least three samples. At the sample level, PCA and Cook’s distance were used to identify potential outliers. Both approaches identified one 17-month male sample as a source of outlier expression, and this sample also had the lowest RIN (7.0). With this outlier sample removed, isoform-level abundances of the remaining 16 samples were analyzed with IsoformSwitchAnalyzeR (v2.1.2, R v4.2). Isoform fractions (IF) were calculated independently for each age group. Counts assigned to each *Bdnf* isoform were divided by the sum of all *Bdnf* counts across all samples of that age.

#### Restraint stress model

Adult male mice were divided into two cohorts, control and stressed. Mice in stressed cohorts were restrained for 1 hour every day for 5 consecutive days and were then released back into their cages. The timing of the restraint during the day was varied across the 5 days to reduce habituation. By contrast, the mice in the control group were handled similarly, but were not restrained and were returned to the cage immediately after handling. The whole gut transit time (WGTT) of the mice in response to treatment was assayed on the day after the last day of restraint. In a separate cohort of mice, 2 hours after the last restraint stress, the animals were sacrificed, and their small intestinal LM-MP tissues were isolated and snap frozen to analyze changes in expression of various *Bdnf* isoform transcripts.

#### Dexamethasone induced model of GI dysmotility

Adult male mice were divided into two cohorts, control and DEXA. Mice in the control group were treated with vehicle (10% DMSO in saline, intraperitoneal route) and those in the DEXA group were treated with dexamethasone (MedChemExpress, Cat. No. HY-14648; 5mg/kg body weight in 10% DMSO, intraperitoneal route). Dexamethasone is light sensitive so was shielded from bright light, both in powder form and in solution. Mice were first housed in individual cages and after a habituation phase lasting 30min, they were treated with vehicle or dexamethasone, which were administered intraperitoneally using a sterile 1 ml insulin syringe. The mice were subjected to WGTT assay 30 minutes after the treatment.

Another 2 cohorts of mice were similarly used, wherein the control cohort was dosed similarly with vehicle and the DEXA group was dosed similarly with dexamethasone. 4 hours after the treatment, the mice were euthanized, and their small intestinal LM-MP tissues were isolated and immediately snap-frozen in liquid nitrogen.

#### Effect of TrkB agonist HIOC on dexamethasone-induced model of GI dysmotility

HIOC (N-[2-(5-hydroxy-1H-indol-3-yl)ethyl]-2-oxopiperidine-3-carboxamide) is a known specific agonist of TrkB. Adult male mice were divided into two cohorts, DEXA and DEXA+HIOC. Mice were first housed in individual cages and then those in the DEXA+HIOC group were treated with HIOC (MedChemExpress, Cat. No. HY-101446; 50 mg/kg body weight in 10% DMSO, intraperitoneal route), while those in DEXA group were treated with Vehicle (10% DMSO in saline, intraperitoneal route). The mice were then released back to their individual cages for 30 minutes after which both groups were treated with dexamethasone (5 mg/kg body weight in 10% DMSO, intraperitoneal route) and again released back to their individual cages for an additional 30 minutes. Treatment was administered intraperitoneally using a sterile 1 ml insulin syringe. The mice were subjected to whole gut transit time assays 30 minutes after the dexamethasone treatment.

#### Whole gut transit time analyses (WGTT)

These experiments were started between the times of 8 and 9 AM. Mice were individually caged in plastic cages without bedding and were left undisturbed 30 minutes. Water was provided to the mice before and during the experiment *ad libitum*. All mice received oral gavage of 300 μl of 6% (w/v) carmine red (Sigma C1022) in 0.5% (w/v) methylcellulose (Sigma M0512) in sterile saline. The oral gavage was performed such that the entirety of the dye suspension was delivered into the animal’s stomach. The mice were then left undisturbed in their individual cages for 70 minutes after which their cages were checked every 15 minutes for the presence of red colored fecal pellets. The time post-gavage for every mouse to produce a red colored fecal pellet was measured, and the difference between the gavage time and red fecal pellet production time was established as the whole gut transit time for that mouse. The experiment was terminated at 250 minutes post-gavage and the WGTT of any mice that did not expel the red dye at the termination was marked at the value of 265 (i.e. 250 + 15) min. The mean difference in WGTT (in minutes) between cohorts was analyzed statistically.

#### Mouse small intestine LM-MP tissue isolation for non-sequencing experiments

Mice were anesthetized with isoflurane and sacrificed by cervical dislocation. Mice were placed in dorsal recumbency on the surgical surface and the skin was disinfected with 70% EtOH before opening the abdominal cavity and collecting the small intestine (SI) into a petri dish containing PBS with 1x Pen-Strep. Sterile PBS solution was flushed through the intestine using a 20ml syringe to remove fecal matter. Entire small intestine was cut into 2cm pieces into a sterile petri dish containing Opti-MEM medium with 1X Pen-Strep. SI segments were placed over a sterile 1ml pipette and LM-MP tissue was peeled off from the underlying tissue using a wet sterile cotton swab. Tissues isolated for qPCR-based assessment of *Bdnf* levels were snap frozen immediately after isolation. Tissues isolated for culture were placed in a fresh sterile ice-cold Opti-MEM medium for further use. Tissues isolated for immunostaining experiments were flattened, fixed in freshly prepared ice-cold 4% paraformaldehyde solution for 5 minutes (using our established protocol (Gorecki *et al*, 2025; Kulkarni *et al*, 2023)) and then stored in 1X sterile PBS for subsequent processing.

#### *In vitro* LM-MP tissue culture and treatments

Isolated LM-MP tissues were incubated in standard tissue culture conditions of 37°C and 5% CO_2_, in a sterile 24 well dish containing Stem cell media (SCM, made up of Neurobasal medium containing L-glutamine, B27, and Pen-Strep (all Invitrogen) and Bovine Serum Albumin (Sigma) (Kulkarni *et al*, 2017)) and treated with Vehicle (10% DMSO in saline) or dexamethasone (5µM in 10% DMSO) for 4 hrs. After the incubation, tissues were snap frozen in liquid nitrogen & stored at −80°C for further experimental analysis. To assess how soon after dexamethasone exposure does the expression of *Bdnf IV* and *VI* isoforms significantly vary, we cultured freshly isolated LM-MP in SCM that either contained dexamethasone (5µM in 10% DMSO) or vehicle (10% DMSO) for 1-hour or 2-hours. Immediately after the culture, the tissues were snap frozen and used for subsequent RNA-based analyses.

#### RNA isolation and quantitative real-time PCR

Total RNA was extracted from frozen LM-MP tissues with Direct-zol RNA miniprep kit (Zymo research) and 1 μg RNA was converted to cDNA using SuperScript III (Thermo, Cat.No.11752050) in a 20µl reaction. Quantitative real-time PCR was performed on a QuantStudio3 real time PCR system (Life Technologies, USA) with TaqMan Fast advanced qPCR Master Mix (Cat. No. 4444557, Thermo Scientific, USA) using gene specific TaqMan assay probes. Mouse *Hprt* served as a housekeeping gene for normalization. Primer and probe information is provided in Table 1. 20µl qRT-PCR reactions were incubated at 95℃ for 10 min, followed by 40 cycles at 95 ℃ for 10 s, 60℃ for 10 s, and 72 ℃ for 40 s. Relative fold changes of genes between treatment groups were calculated using the 2^−ΔΔCt^ method. Primer information is provided in Table 1, and primers specific to TrkB.FL and T1 were obtained from Tessarollo Lab (Tomassoni-Ardori *et al*, 2019).

#### Protein isolation and ELISA

Proteins from frozen LM-MP were extracted using a cell lysis buffer containing RIPA buffer (150 mM NaCl, 50 mM Tris-HCl pH 8.0, 1% NP-40, 0.1% sodium deoxycholate and 0.1% SDS, adjusted with distilled water). Succinctly, for every LM-MP peel, we add 20 μl of 1X RIPA+++ buffer which contains RIPA with Halt Protease Inhibitor (Invitrogen, Cat. No. 78438) at a conc. of 5-10X and Phosphatase Inhibitors 2 and 3 (Sigma Aldrich, Cat. Nos. P5726 and P0044) at a concentration of 2-3% to locking sterile Eppendorf tubes and then add 6-8 sterile autoclaved silicon beads. Tissue was homogenized using a bead beater in cold room at 7-8 speed setting for 5 mins, after which the tubes were shaken at 200-250 rpm for 30-40 mins in cold room, and centrifuged at over 12,000rpm for 20 mins at 4°C. Supernatant thus generated was collected to be used for ELISA. The levels of BDNF in the LM-MP tissues cultured with and without dexamethasone were measured using the BDNF DuoSet kit (R&D Systems; DY248). In brief, sterile and clean ELISA-grade 96-well plates (Corning) were coated with 100 μl of diluted capture antibody/well and incubated overnight at room temperature. On the second day, the plates were washed and blocked with reagent diluent (300 μl for 1 hour at room temperature and 100 μl of protein extract, standards in reagent diluent, or appropriate diluent were added to the prepared wells and incubated for 2 hours at room temperature. Following this, 100 μl each of detection antibody, working dilution of streptavidin-HRP, substrate solution, and 50 μl of stop solution were sequentially added to each well. The optical density of each well was measured using a microplate reader set to 540 nm. The BDNF concentration was expressed in picograms of BDNF per microgram of protein.

#### Immunostaining and imaging murine small intestinal LM-MP tissues

Fixed LM-MP tissues were blocked and permeabilized with 5% normal goat serum and 0.5% Triton-X in 1X sterile PBS for 30 minutes at room temperature. For assessing the co-expression of glucocorticoid receptor GR with all enteric neurons, we performed immunostaining of adult C57BL/6 NIA mice with anti-GR (*Nr3c1*) antibodies (Cell Signaling: CST3660T, 1:1000) and ANNA-1 patient serum containing anti-Hu antibodies (1:1000). For assessing proportions of BDNF-expressing and TrkB-expressing neurons, we immunostained LM-MP tissues from TrkB-GFP mice with antibodies against BDNF (abcam, 1:500), against GFP (Aves Labs, GFP1020, 1:750), and against Hu using ANNA-1 antiserum containing anti-Hu antibodies. After counterstaining with appropriate secondary antibodies and nuclear dye DAPI, the tissues were mounted with Prolong Gold Anti-fade mountant (Invitrogen) and imaged under Leica Stellaris Confocal microscope using 63X PL APO oil immersion objective (NA: 1.40) with z-step set at 0.85 µm. The images were analyzed with Fiji.

Statistics: Statistical analysis to assess differences between group means was conducted using Student’s t-test in Prism 10 software.

## Results

### Restraint stress in adult mice slows down whole gut motility

We applied restraint stress daily for 1 hour for 5 consecutive days on a cohort of adult male C57BL/6 mice and found that restraint stress caused a significant slowing of whole GI motility, as assayed by the carmine-red dye method (n = 5/cohort; mean ± S.E. of WGTT (in min): Control: 93.0 ± 5.612; Stress: 123.0 ± 8.746, p = 0.02, Students’ t-test; **Fig 1**). This replicates a prior study which reports a similar effect of physiological stress on whole GI motility (Schneider *et al*., 2023)

**Figure 1.**
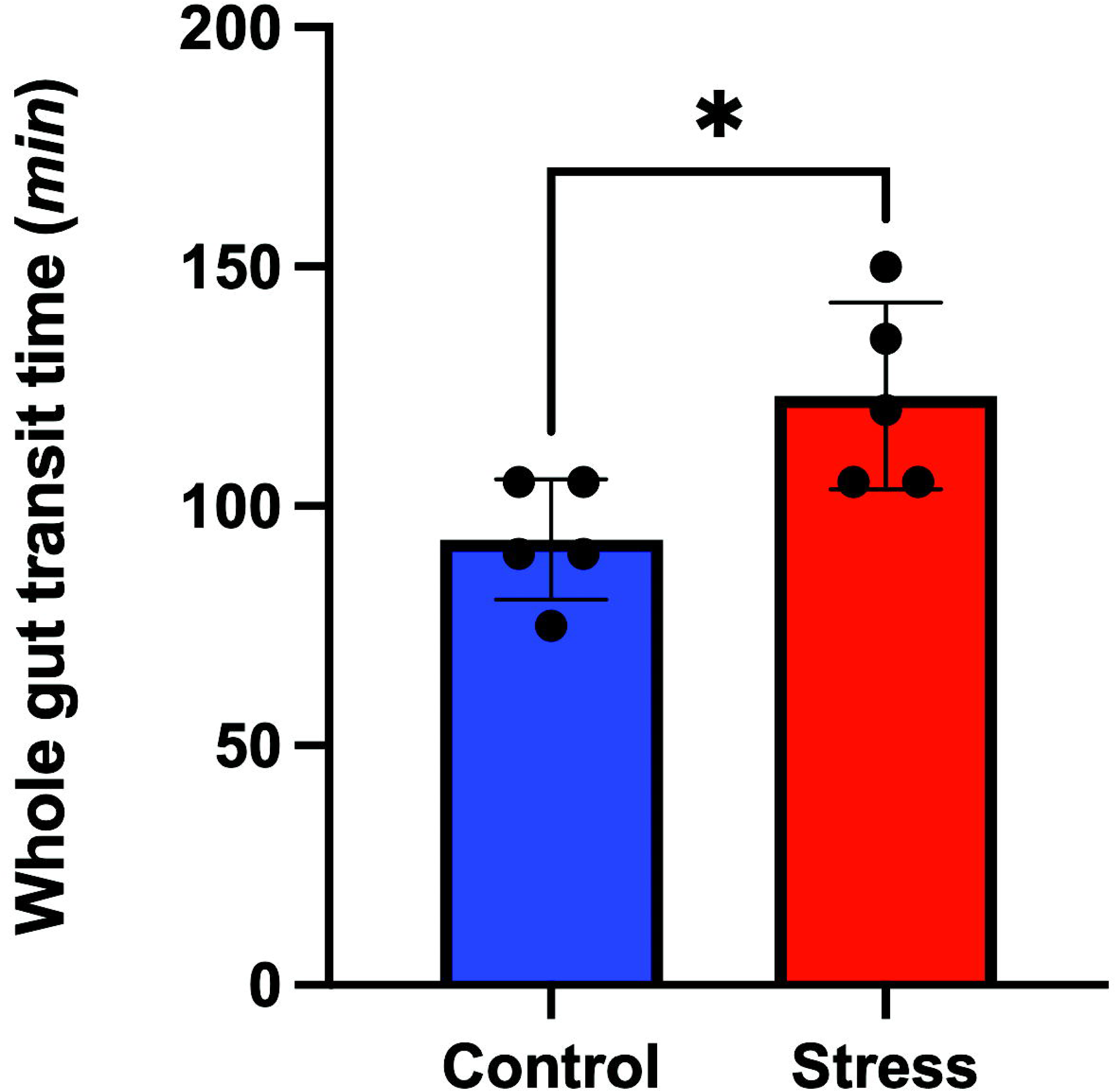
Restraint stress significantly increases whole gut transit time in adult male mice. Graphical representation of the mean ± standard error of the whole gut transit times (assayed using the carmine-red dye method) of two cohorts of mice, where the mice in the ‘Stress’ cohort were restrained for an hour every day for 5 days, and the ‘Control’ cohort mice were handled similarly but not restrained. Mice in the Stress cohort showed significantly longer whole gut transit times on average than mice in the control cohort (n = 5/cohort; mean ± S.E. of WGTT (in min): Control: 93.0 ± 5.612; Stress: 123.0 ± 8.746, p = 0.02, Students’ t-test *p < 0.05. Student’s t-test).

### Stress-responsive *Bdnf* isoforms are expressed in the murine longitudinal muscle - myenteric plexus tissue

Having confirmed that restraint stress delays whole GI transit, we next tested whether stress-responsive *Bdnf* isoforms are detected in the gut wall. For this, we characterized the expression profile of *Bdnf* in the motility-regulating longitudinal muscle and myenteric plexus (LM-MP) tissue of post-natal gut across three ages. We isolated LM-MP tissue from the ileum of juvenile (1 month), adult (6 month), and aging (17 month) male and female mice (n = 3/sex for 1-month and 6-month-old mice, n = 2 males and 2 females for 17-month-old mice) and performed bulk RNA sequencing. First, we confirmed that *Bdnf* transcripts are detected in the ileal tissues of male and female mice at all ages. We next analyzed whether the stress-responsive *Bdnf IV* (from e4) and *Bdnf VI* (from e6) are expressed in the ileal LM-MP tissue across ages and in the two sexes. We found that at all profiled ages, *Bdnf VI* is the dominant isoform, making up 85.3% (95% CI [81.5%, 89.0%]) of all detected *Bdnf* transcripts (**Fig 2**). Similarly, *Bdnf V* (from e5) expression is detected at all ages and in both male and female tissues (percent representation of *Bdnf V* at 1-month: 9.7%, 6-month: 9.4%, and 17-month: 6.2%). By contrast, the other stress-responsive isoform *Bdnf IV* is detected at 1-month and 6-months of age in both male and female ileal LM-MP (percent representation of *Bdnf IV* at 1-month: 5.3%, and 6-month: 2.3%), but it is not detected in the aging ileal LM-MP – suggesting that the expression of this isoform is reduced or lost with aging. On the other hand, the expression of *Bdnf II* (e2) is not detected at 1-month of age but is detected at 6-months and 17-months of age (percent representation of *Bdnf II* at 6-month: 3.3%, and 17-month: 5.1%). 6-months age is the only age we queried in the bulk RNA-seq where we detected the expression of *Bdnf-III* in the murine ileal LM-MP (representing 2.9% of all *Bdnf* transcripts at this age) (**Extended Fig 2-1**). These data show that stress-responsive *Bdnf* isoforms (*IV* and *VI*) represent 85% of all *Bdnf* transcripts across age in the murine ileal LM-MP tissue.

**Figure 2.**
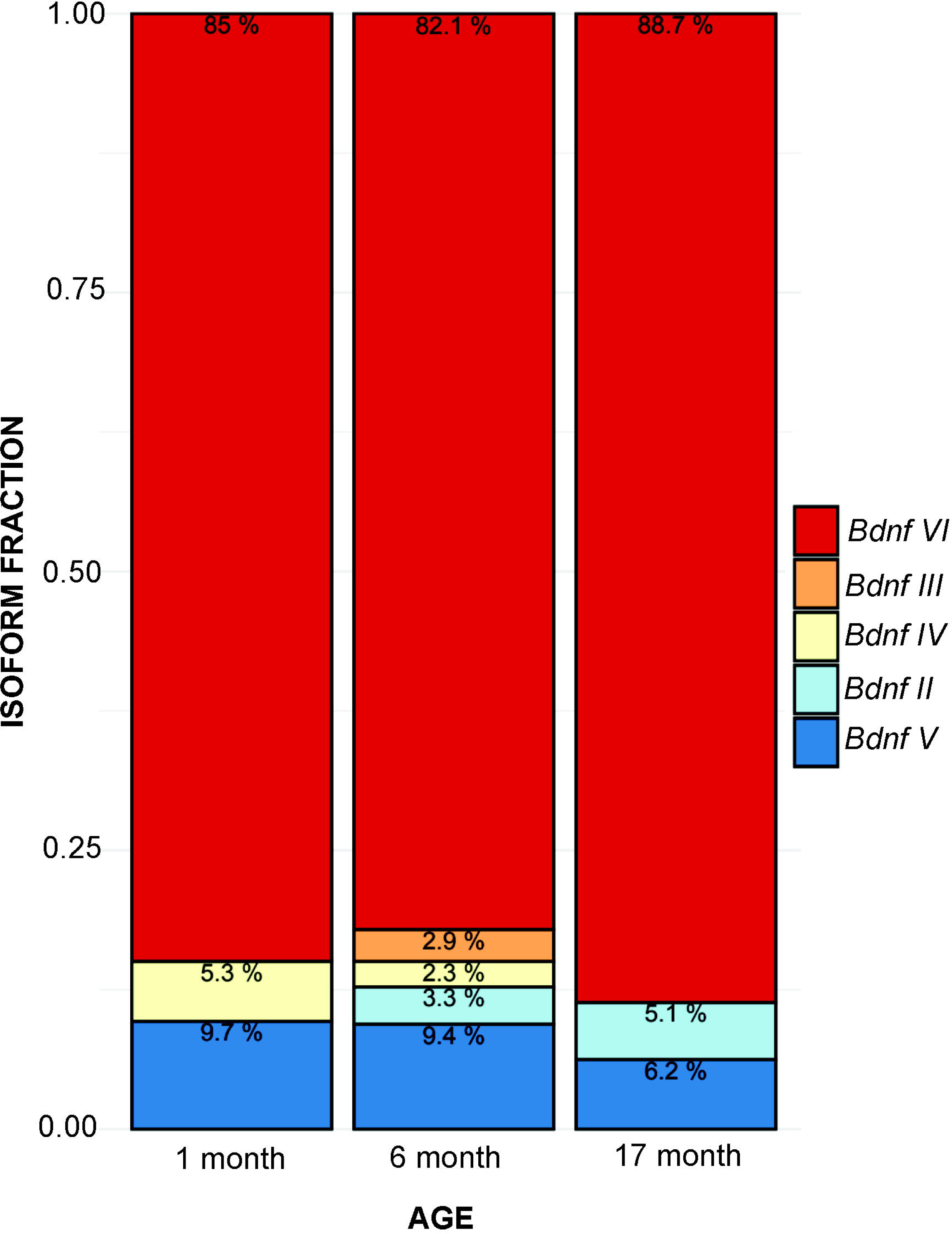
*Bdnf-IV and VI* together account for >85% of all *Bdnf* isoforms in the ileal LM-MP at all post-natal ages. Graphical representation of the proportions of various *Bdnf* isoforms whose transcripts were detected in the murine ileal LM-MP layer in two different sexes across three different ages of 1-month (juvenile), 6-month (mature adult), and 17-month (aging). This representation shows that stress-responsive *Bdnf-VI* is the dominant isoform found in the murine ileal LM-MP at all ages, with its proportions accounting for >80% of all isoforms. The isoform *Bdnf-V* is the second most represented isoform at all ages in the murine LM-MP layer, but little is known about the mechanisms that regulate its expression. In addition to *Bdnf-VI, Bdnf-IV* is another stress-responsive isoform detected in the juvenile and adult mice but is not detected in aging murine ileal LM-MP. In contrast to *Bdnf-IV, Bdnf-II* isoform is detected in the adult and aging ileal tissues, but not in the juvenile tissues. Finally, *Bdnf-III* isoform is detected only in the adult ileal LM-MP and not at the other ages. Detailed age-specific isoform specific abundance of various Bdnf isoforms presented in Extended Figure 2 – 1.

### BDNF and TrkB are expressed by myenteric neurons

Prior studies have used immunohistochemical and conditional transgenic mice to establish the expression of BDNF in myenteric ganglia in the post-natal gut (Kovler *et al*, 2021; Singh *et al*, 2022). However, the exact nature and proportions of adult small intestinal myenteric cells that express BDNF have not been determined. By contrast, using a reporter transgenic mouse, it has been demonstrated that TrkB is expressed by myenteric neurons in the adult gut (Nestor-Kalinoski *et al*., 2022). To support this result, we performed immunohistochemistry with BDNF and Hu antibodies using TrkB-GFP mice to confirm that BDNF and TrkB are expressed by enteric neurons. We observed that BDNF and TrkB are expressed exclusively by enteric neurons (**Fig 3A**). On quantification of 448 Hu-expressing cells from 3 mice, we found that ∼24% of all Hu-expressing myenteric cells were immunoreactive for BDNF (average ± standard error of percent of all Hu^+^ cells immunoreactive for BDNF: 23.65 ± 0.51) while a non-overlapping ∼16% of all Hu-expressing myenteric cells expressed TrkB (average ± standard error of percent of all Hu^+^ cells expressing TrkB-GFP: 16.09 ± 1.87) (**Fig 3B**). This data shows that BDNF and TrkB are neuronally expressed genes in the myenteric plexus. We further performed immunohistochemistry using anti-BDNF antibodies on formalin fixed paraffin embedded (FFPE) tissue sections from the adult small intestine of human patients without any known motility disorder and again found that while BDNF protein was present in the myenteric ganglia, its abundance was significantly higher in neuron-like cells (**Fig 3C**).

**Figure 3.**
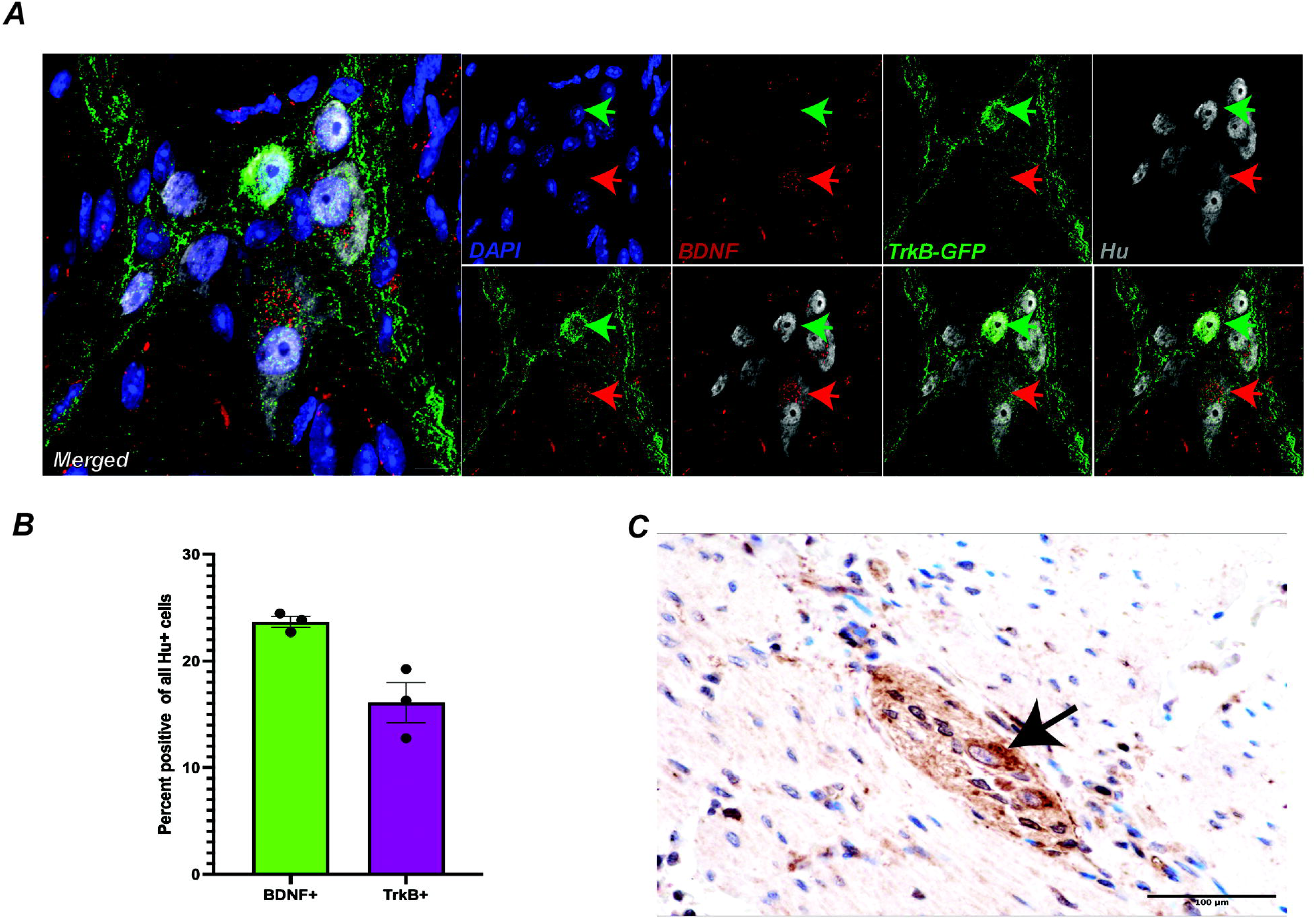
*Bdnf* is a neuronally expressed gene in the adult murine small intestinal ENS. Transcriptomic and proteomic evidence of neuronal expression of BDNF. (A) 3D projection image obtained by confocal microscopy of a representative murine adult small intestinal myenteric ganglion from a TrkB-GFP mouse, where GFP expression (green) is enhanced by immunostaining with anti-GFP antibodies, and antibodies against BDNF (red) and ANNA-1 antisera containing anti-Hu antibodies (grey) label BDNF-expressing cells and Hu-expressing myenteric neurons, respectively. Immunolabeling shows that TrkB-GFP-expressing cells (green arrow) and BDNF-expressing cells (red arrow) are both myenteric neurons. Nuclei are labeled with DAPI (blue) and scale bar represents 10 μm. (B) Quantification of BDNF-immunoreactive and TrkB-GFP^+^ neurons in the small intestinal myenteric plexus tissue of 3 adult mice. (C) Immunostaining a human small intestinal tissue section with antibodies against BDNF also shows that while the secreted form of BDNF is present inside and outside of the myenteric ganglia, the highly enriched presence of BDNF in the cytoplasm of a myenteric neuron (black arrow) suggests its likely source of expression. Scale bar represents 100 μm.

### Restraint stress causes a significant reduction in *Bdnf IV* expression

Since stress has been observed to reduce *Bdnf* expression in the central nervous system (CNS) (Smith *et al*, 1995), we next tested whether restraint stress-induced slowdown of GI motility is associated with alterations in expression of different *Bdnf* isoforms. Using LM-MP tissues derived from restrained (i.e. stressed) and restraint-naïve control mice (n = 5/cohort) and by using quantitative real time PCR (qRT-PCR) with isoform-specific primers, we found that 5 days of daily restraint stress caused a significant reduction in the expression of stress-responsive *Bdnf IV* isoform compared to control mice (n = 5/cohort; mean ± S.E. fold change of *Bdnf IV*: Control: 1.002 ± 0.034; Stress: 0.800 ± 0.044, p = 0.006; Students’ t-test) while none of the other tested *Bdnf* isoforms (*I, II, III, V* or *VI*) showed any significant change (**Fig 4**).

**Figure 4:**
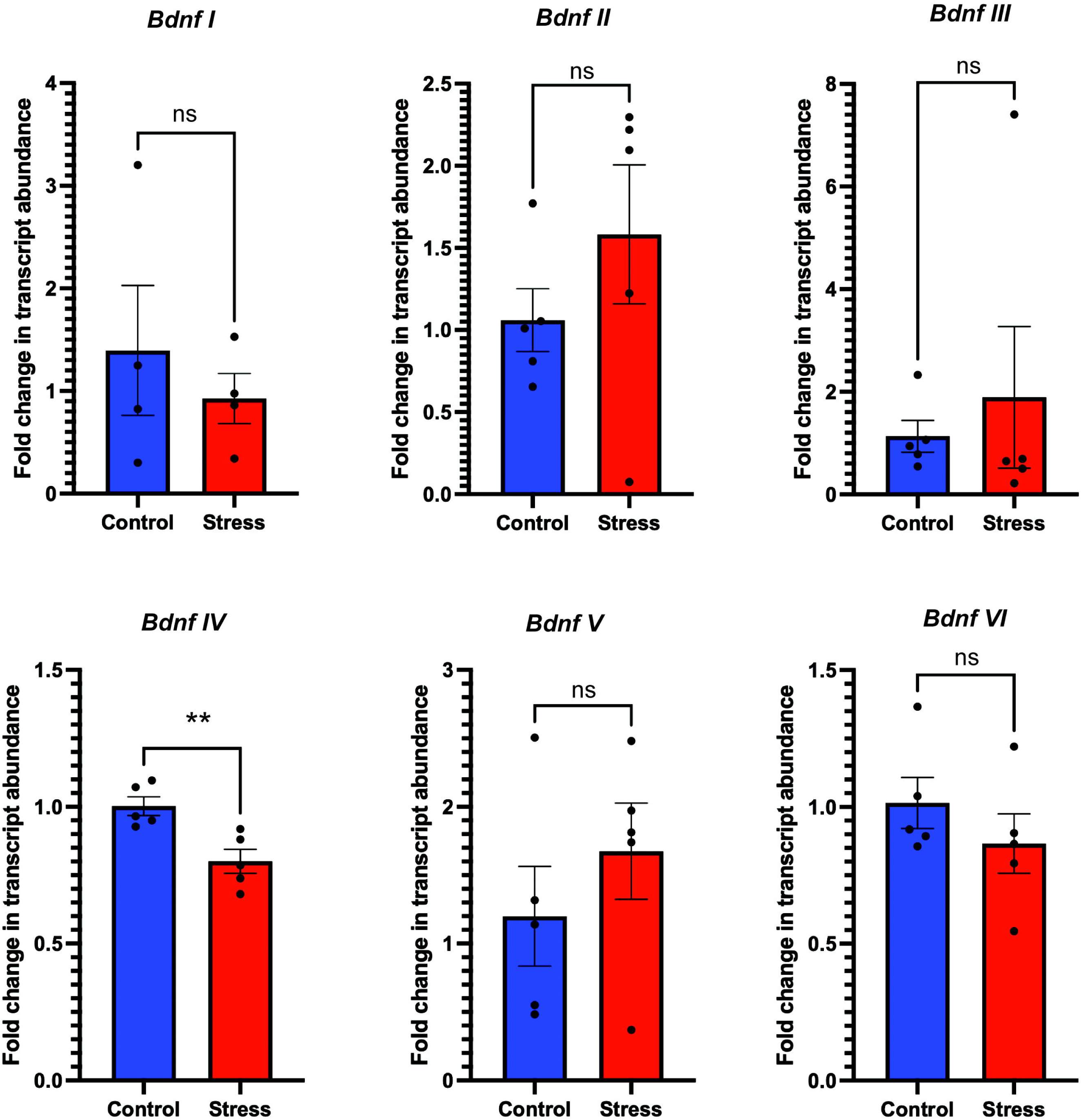
Restraint stress causes a significant reduction in *Bdnf-IV* expression. Graphical representations of the mean ± standard error of fold change in transcript abundance of selected *Bdnf* isoforms (calculated using the 2^−ΔΔCt^ method) in small intestinal LM-MP tissues derived from two cohorts of mice – relative to the expression of housekeeping gene *Hprt*. Mice in the ‘Stress’ cohort were restrained for an hour every day for 5 days, and the ‘Control’ cohort mice were handled similarly but not restrained. Mice in the Stress cohort showed significantly reduced expression of *Bdnf-IV* isoform while no other isoform showed significant reduction in expression compared to control cohort (n = 5/cohort; mean ± S.E. fold change of *Bdnf-IV*: Control: 1.002 ± 0.034; Stress: 0.800 ± 0.044, p = 0.006; Students’ t-test).

### Glucocorticoid receptor is expressed by all cells of the myenteric ganglion

The Thaiss lab showed that stress significantly increases serum corticosterone levels (Schneider *et al*., 2023). To study whether the increased endogenous glucocorticoid has a direct effect on myenteric neurons, we first tested whether glucocorticoid receptor GR is expressed by myenteric neurons. By performing immunostaining on small intestinal myenteric plexus tissue with anti-GR antibodies and ANNA-1 antisera, we observed that GR is expressed ubiquitously by all LM-MP cells, including Hu-expressing cells of the myenteric ganglia (**Fig 5A**). This suggests that stress-mediated increase in corticosterone can directly influence all neurons, including *Bdnf*-expressing myenteric neurons, through their expression of GR.

**Figure 5:**
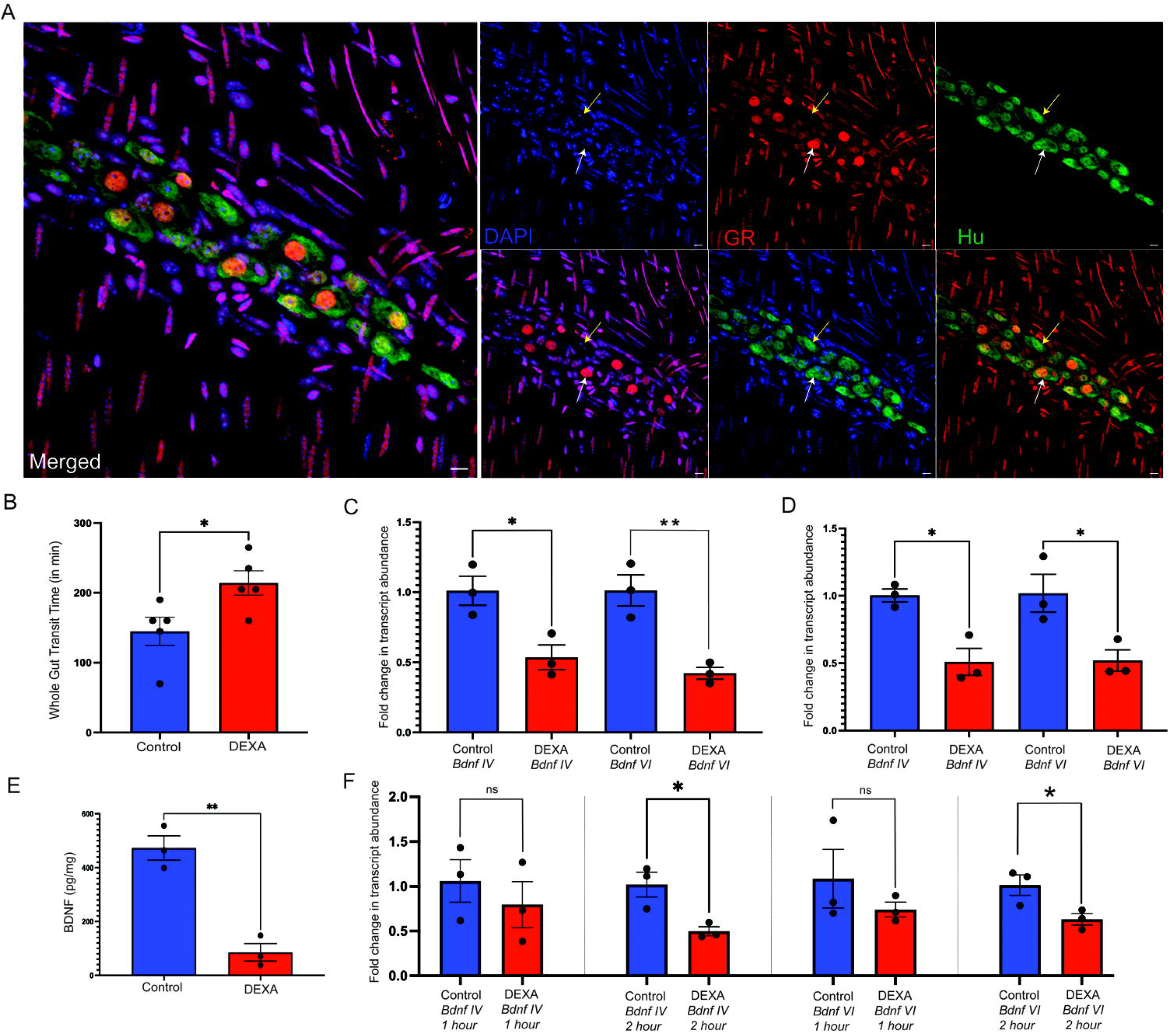
Dexamethasone exposure significantly delays whole gut transit while downregulating myenteric BDNF expression. (A) Representative of a myenteric ganglion and surrounding cells from an adult murine small intestinal LM-MP tissue when immunostained with antibodies against glucocorticoid receptor (GR; red) and with ANNA-1 antisera containing anti-Hu antibodies (green) shows that GR is expressed by all cells of the LM-MP tissues. GR shows strong nuclear immunoreactivity in a subset of Hu-expressing myenteric cells (white arrow) while showing relatively weaker nuclear immunoreactivity in other cells (yellow arrow). Nuclei are stained with DAPI (blue). Scale bar is 10 μm. (B) Graphical representation of the mean ± standard error of the whole gut transit times (assayed using the carmine-red dye method) of two cohorts of mice, where the mice in the ‘DEXA’ cohort were dosed once with dexamethasone (5mg/Kg body weight), and the ‘Control’ cohort mice were dosed with Vehicle (10% DMSO in Saline). Mice in the DEXA cohort showed significantly longer whole gut transit times on average than mice in the control cohort (n = 5/cohort; mean ± S.E. of WGTT (in min): Control: 145.0 ± 20.12; DEXA: 214.0 ± 17.49, p = 0.03, Students’ t-test). (C) Adult male mice dosed with dexamethasone (DEXA) and sacrificed 4 hours later showed a significant down regulation in the expression of stress and glucocorticoid-responsive *Bdnf IV* and *Bdnf VI* transcripts in their small intestinal LM-MP tissues, when compared to age- and sex-matched mice that were dosed with vehicle (n = 3/cohort; mean ± S.E. fold change of *Bdnf IV*: Control: 1.011 ± 0.104; DEXA: 0.536 ± 0.087, p = 0.025; mean ± S.E. fold change of *Bdnf VI*: Control: 1.012 ± 0.110; DEXA: 0.422 ± 0.042, p = 0.007; Students’ t-test). (D) Small intestinal LM-MP tissues from adult male mice that were cultured with dexamethasone for 4 hours showed a significant reduction in the abundance of both *Bdnf IV* and *Bdnf VI* transcripts (n = 3/cohort; mean ± S.E. fold change of *Bdnf IV*: Control: 1.002 ± 0.047; DEXA: 0.510 ± 0.099, p = 0.030; mean ± S.E. fold change of *Bdnf VI*: Control: 1.018 ± 0.140; DEXA: 0.520 ± 0.079, p = 0.028; Students’ t-test). (E) Small intestinal LM-MP tissues from adult male mice that were cultured with dexamethasone for 4 hours showed a significant reduction in the abundance of BDNF protein as assessed by ELISA, when compared to tissues from age- and sex-matched mice that were cultured with vehicle. **p < 0.01, Students’ t-test (n = 3/cohort; mean ± S.E. BDNF levels (in pg/g of tissue): Control: 473.0 ± 45.32; DEXA: 85.5 ± 32.63, p = 0.002; Students’ t-test). (F) Small intestinal LM-MP tissues from adult male mice that were cultured with dexamethasone for 1 hour or for 2 hours showed that while 1 hour of exposure to dexamethasone caused no significant change in the expression of either *Bdnf IV* or *VI* transcripts, continued exposure of LM-MP to dexamethasone for 2 hours caused a significant reduction in the abundance of both *Bdnf IV* and *Bdnf VI* transcripts n = 3/treatment at 2 hours; mean ± S.E. fold change of *Bdnf-IV*: Control: 1.020 ± 0.137; DEXA: 0.497 ± 0.049, p = 0.023; mean ± S.E. fold change of *Bdnf-VI*: Control: 1.014 ± 0.114; DEXA: 0.630 ± 0.063, p = 0.043; Students’ t-test).

### Dexamethasone exposure significantly delays whole gut transit time while downregulating myenteric BDNF expression

To test whether GR stimulation alone, without restraint stress, similarly alters GI motility, we tested the effect of dexamethasone, a synthetic fluorinated homologue of hydrocortisone and a mimetic of endogenous glucocorticoid (Dubashynskaya *et al*, 2021), on whole gut transit time (WGTT). Mice treated with dexamethasone showed a significant delay in their whole GI motility, when compared to vehicle-treated control mice (n = 5/cohort; mean ± S.E. of WGTT (in min): Control: 145.0 ± 20.12; DEXA: 214.0 ± 17.49, p = 0.03, Students’ t-test; **Fig 5B**). This replicates an independent finding that observed significant delay in whole gut transit in dexamethasone-treated mice, albeit after multiple doses of dexamethasone (Schneider *et al*., 2023).

We next asked if *Bdnf* expression is altered by dexamethasone exposure, similarly to the effect of restraint-stress. For this, we tested whether exposure to dexamethasone causes a significant reduction in the expression of stress-responsive *Bdnf IV* and/or *VI* isoforms. In small intestinal LM-MP of adult male mice treated with dexamethasone (DEXA cohort) or vehicle (control cohort) *in vivo*, we assessed the abundance of these *Bdnf* isoforms at 4 hours post-treatment, a timepoint aligning with the duration of our WGTT experiments. By performing qRT-PCR, we found that expression of both *Bdnf IV* and *VI* were significantly reduced in small intestinal LM-MP of mice treated with dexamethasone when compared to vehicle treated mice in the control cohort (n = 3/cohort; mean ± S.E. fold change of *Bdnf IV*: Control: 1.011 ± 0.104; DEXA: 0.536 ± 0.087, p = 0.025; mean ± S.E. fold change of *Bdnf VI*: Control: 1.012 ± 0.110; DEXA: 0.422 ± 0.042, p = 0.007; Students’ t-test; **Fig 5C**).

*Bdnf* is an immediate early gene whose expression may be triggered by tissue damage (Ballarín *et al*, 1991; Hughes *et al*, 1993). To ensure that our results are not affected by the dissection of tissues immediately before RNA isolation, we cultured adult small intestinal LM-MP tissues from male mice in stem cell media with vehicle (DMSO) or dexamethasone (5 µM) for 4 hours and performed qRT-PCR with *Bdnf* isoform-specific probes. Again, we found that exposure to dexamethasone significantly reduced the expression of both stress-responsive isoforms *Bdnf IV* and *Bdnf VI* when compared to control tissues that were similarly cultured but were instead exposed to vehicle (n = 3/cohort; mean ± S.E. fold change of *Bdnf IV*: Control: 1.002 ± 0.047; DEXA: 0.510 ± 0.099, p = 0.030; mean ± S.E. fold change of *Bdnf VI*: Control: 1.018 ± 0.140; DEXA: 0.520 ± 0.079, p = 0.028; Students’ t-test; **Fig 5D**).

Next, we used the same culture system with and without dexamethasone for assessing whether exposure to dexamethasone alters BDNF protein levels in LM-MP tissue. By performing an ELISA, we found that LM-MP cultured with dexamethasone for 4 hours showed significant reduction in the levels of BDNF protein when compared to LM-MP cultured with vehicle (n = 3/cohort; mean ± S.E. BDNF levels (in pg/g of tissue): Control: 473.0 ± 45.32; DEXA: 85.5 ± 32.63, p = 0.002; Students’ t-test; **Fig 5E**).

Having observed that dexamethasone exposure suppresses *Bdnf* expression in LM-MP tissue at 4 hours—the duration of the whole gut transit assay—we next sought to identify the earliest timepoint at which continued dexamethasone exposure significantly alters *Bdnf* transcript levels. We cultured LM-MP tissues in media containing vehicle (control cohort) or dexamethasone (DEXA cohort) for 1 hour or 2 hours and assessed the transcript abundance of *Bdnf-IV* and *VI* at these timepoints when compared to control tissues that were similarly cultured but were instead exposed to vehicle. We found that while there was no significant change in the expression of either *Bdnf* isoforms at the 1 hour timepoint, expression of both *Bdnf-IV* and *VI* isoforms were significantly reduced at the 2 hour timepoint (n = 3/treatment at 2 hours; mean ± S.E. fold change of *Bdnf-IV*: Control: 1.020 ± 0.137; DEXA: 0.497 ± 0.049, p = 0.023; mean ± S.E. fold change of *Bdnf-VI*: Control: 1.014 ± 0.114; DEXA: 0.630 ± 0.063, p = 0.043; Students’ t-test; **Fig 5F**). Given that whole gut transit time experiments were started 30 minutes after dexamethasone injection, this would suggest that dexamethasone-mediated alterations to *Bdnf-IV* and *VI* expression occur between the first 30 – 90 minutes of the 4-hour experiment.

### Deletion of TrkB in enteric neurons delays motility

We observed that DEXA and stress-induced loss of BDNF is associated with GI dysmotility, and prior data has shown using the hemizygous *Bdnf* ^+/-^ mutant mice that reduced BDNF also drives GI dysmotility (Chen *et al*., 2014). We next aimed to investigate how loss of TrkB in ENS affects GI motility. Alternate splicing in the Ntrk2 gene produces two TrkB isoforms: A full-length isoform (TrkB.FL) that contains extracellular and intracellular domains and hence carries out intracellular cell signaling in response to extracellular BDNF, or a truncated (TrkB.T1) isoform that lacks the intracellular kinase domain and thus acts as a sink for BDNF (Biffo *et al*, 1995; Eide *et al*, 1996). To understand the native proportion of these isoformsn in the adult murine ENS, we first queried their relative expression using isoform specific primers (Table 1). Here, we observed that at steady-state, expression of both FL and T1 isoforms is detected in the adult murine small intestinal LM-MP tissue, with TrkB.T1 having significantly greater abundance than the TrkB.FL isoform (n = 3; mean ± S.E. Delta Ct of TrkB.FL: 8.653 ± 0.1771; Delta Ct of TrkB.T1: 5.882 ± 0.5381; p = 0.008; Students’ t-test **Fig 6A**).

**Figure 6:**
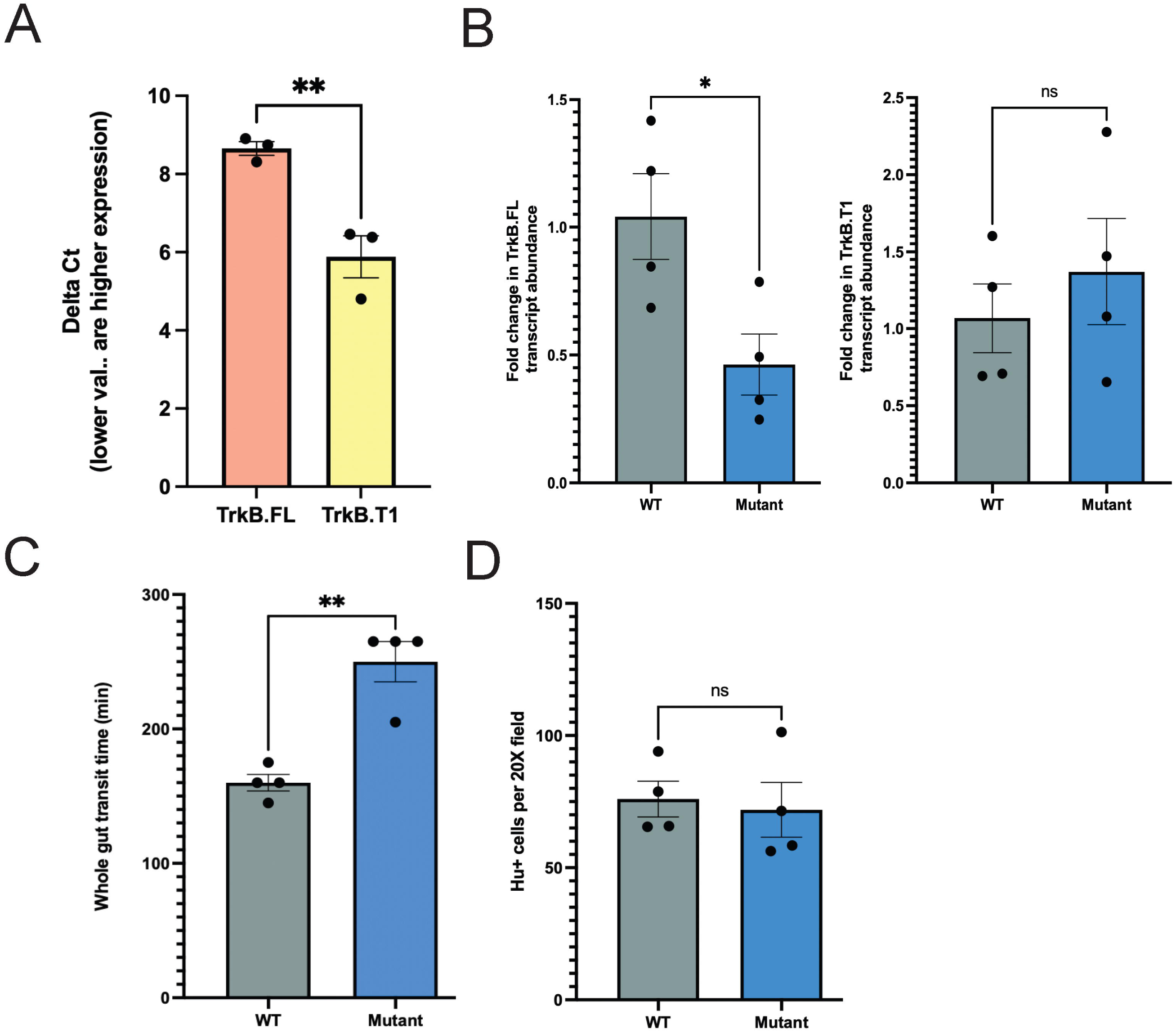
Conditional mutation of *Ntrk2* in ENS neurons drives depletion of TrkB.Fl isoform resulting in significant GI dysmotility. (A) Graphical representation of the mean ± standard error of the Delta Ct values of *Ntrk2* transcripts encoding full length TrkB.FL and the truncated TrkB.T1 isoforms in the healthy adult murine small intestinal LM-MP tissues shows that the TrkB.T1-encoding transcripts have significantly lower Delta Ct values (suggesting higher abundance) when compared to the transcripts encoding TrkB.FL isoform (n = 3; mean ± S.E. Delta Ct of TrkB.FL: 8.653 ± 0.1771; Delta Ct of TrkB.T1: 5.882 ± 0.5381; p = 0.008; Students’ t-test). (B) Graphical representation of the mean ± standard error of the fold change of *Ntrk2* tran-scripts encoding full length TrkB.FL and the truncated TrkB.T1 isoforms in the adult murine small intestinal LM-MP tissues of *Mutant Wnt1*-cre:TrkB^fl/fl^ mice compared to *Wnt1*-cre:TrkB^WT/WT^ wildtype (*WT*) control mice, shows a significant decrease in expression of TrkB.FL-encoding transcripts in mutant mice, while level of TrkB.T1-encoding transcripts remained unchanged (n = 4/cohort; mean ± S.E. fold change of TrkB.FL: WT: 1.042 ± 0.1678; mutant: 0.4628 ± 0.1194, p = 0.03; mean ± S.E. fold change of TrkB.T1: WT: 1.069 ± 0.2226; mutant: 1.370 ± 0.3450, p = 0.49; Students’ t-test). (C) Graphical representation of the mean ± standard error of the whole gut transit time (in minutes) of *Mutant Wnt1*-cre:TrkB^fl/fl^ mice and *Wnt1*-cre:TrkB^WT/WT^ wildtype (*WT*) control mice shows that the mutant mice have significantly increased transit time of the carmine red dye – suggesting slower GI motility, when compared to the WT control mice n = 4/cohort; mean ± S.E. of WGTT (in min): WT: 160.0 ± 6.12; mutant: 250.0 ± 15.0, p = 0.001, Students’ t-test). (D) Graphical representation of the mean ± standard error of the density of Hu-immunolabeled cells in the small intestinal LM-MP tissues of *Mutant Wnt1*-cre:TrkB^fl/fl^ mice and *Wnt1*-cre:TrkB^WT/WT^ wildtype (*WT*) control mice shows that the density of Hu^+^ cells in 20X fields remains unchanged in mutant mice when compared to control WT mice.; n = 4/cohort; mean ± S.E. of Hu^+^ cells per 20x field: WT: 75.99 ± 6.759; mutant: 71.89 ± 10.37; p = 0.75, Students’ t-test).

To study the effect of ENS-specific TrkB knockout, we generated *Wnt1*-cre:TrkB^fl/fl^ mice, in which TrkB.FL is homozygously knocked out in *Wnt1*-expressing neural crest and their derivative cells, including in the neural crest-derived cells of the enteric nervous system, through excision of the kinase domain-containing exon 15 (Chen *et al*, 2005; Rios-Pilier & Krimm, 2019). Using this model, we first tested the effect of this conditional TrkB mutation in ENS on the expression of the two TrkB isoforms in the gut wall. Profiling the small intestinal LM-MP of adult *Wnt1*-cre:TrkB^fl/fl^ mutant mice compared to *Wnt1*-cre:TrkB^WT/WT^ wildtype (WT) control mice, we found a significant decrease in expression of transcripts encoding TrkB.FL isoform, while level of transcripts encoding for TrkB.T1 remained unchanged (n = 4/cohort; mean ± S.E. fold change of TrkB.FL: WT: 1.042 ± 0.1678; mutant: 0.4628 ± 0.1194, p = 0.03; mean ± S.E. fold change of TrkB.T1: WT: 1.069 ± 0.2226; mutant: 1.370 ± 0.3450, p = 0.49; Students’ t-test; **Fig 6B**).

Given our observation that dexamethasone exposure both reduces *Bdnf* levels and delays whole gut transit, we hypothesized that loss of TrkB.FL in *Wnt1*-expressing neural crest-derived cells would likewise reduce motility through decreased BDNF-TrkB signaling in enteric neurons. Consistent with this, we observed a significant increase in whole gut transit time (WGTT) in *Wnt1*-cre:TrkB^fl/fl^ mutant mice compared to *Wnt1*-cre:TrkB^WT/WT^ wildtype (WT) control mice (n = 4/cohort; mean ± S.E. of WGTT (in min): WT: 160.0 ± 6.12; mutant: 250.0 ± 15.0, p = 0.001, Students’ t-test; **Fig 6C**).

Finally, we investigated if the conditional *Ntrk2* mutation and resulting significant reduction in TrkB.FL isoform expression and delayed transit time is associated with alterations in the overall neuronal structure of the ENS. For this, we performed immunostaining using antibodies against Hu, which is expressed by myenteric neurons and has been previously used to assess changes to ENS structure(Kulkarni *et al*., 2017), on small intestinal LM-MP from adult *Wnt1*-cre:TrkB^fl/fl^ mutant and *Wnt1*-cre:TrkB^WT/WT^ wildtype (WT) control mice. After imaging 7 – 8 20X fields per mouse and on enumerating of numbers of Hu-immunolabeled cells/20X field, we found no significant difference in cell density of Hu^+^ cells in a 20X field (n = 4/cohort; mean ± S.E. of Hu^+^ cells per 20x field: WT: 75.99 ± 6.759; mutant: 71.89 ± 10.37; p = 0.75, Students’ t-test; **Fig 6D**).

### Targeted activation of TrkB restores motility in dexamethasone-induced delayed transit models

Our observations that reduced *Bdnf* expression is associated with slowed whole gut motility in dexamethasone-treated mice, and that ENS-specific loss of the TrkB.FL isoform, which carries out BDNF signaling, likewise slows motility, suggest that dexamethasone’s effect on whole gut motility may be mediated by a reduction in BDNF–TrkB signaling. This suggests that increasing TrkB signaling would improve the gut motility of dexamethasone-treated mice. To test this hypothesis, we studied whether a synthetic and specific agonist of TrkB can restore whole gut motility of dexamethasone-treated mice. In rodent models, the N-acetyl serotonin (NAS)-derived compound HIOC has been demonstrated to be a specific agonist of TrkB (Dhakal *et al*, 2021). We found that mice pre-treated with HIOC 30 minutes before dexamethasone administration (DEXA + HIOC cohort) had significantly lower whole gut transit times on average than mice treated with dexamethasone alone (DEXA cohort) (n = 9/cohort; mean ± S.E. of WGTT (in min): DEXA: 228.3 ± 7.12; DEXA + HIOC: 150.0 ± 21.94, p = 0.003, Students’ t-test; **Fig 7**).

**Figure 7:**
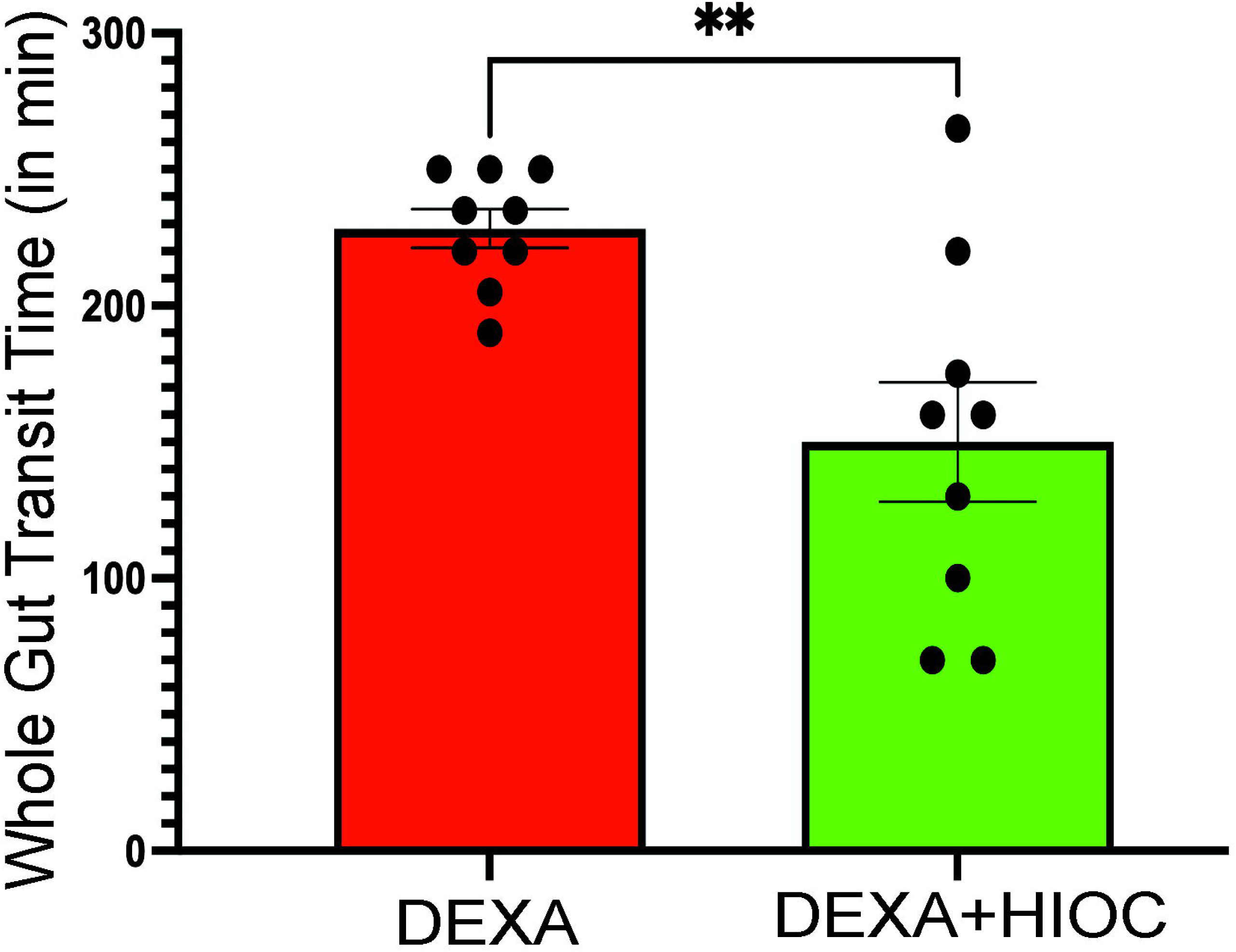
HIOC ameliorates delayed whole gut motility in mice with dexamethasone-induced dysmotility. Dexamethasone-treated adult male mice that were pre-treated with HIOC - the synthetic and specific agonist of TrkB - showed significant reduction in their whole gut transit time (WGTT) of orally gavaged carmine-red dye, and hence improved whole GI motility, when compared to age- and sex-matched mice that were dosed only with dexamethasone (n = 9/cohort; mean ± S.E. of WGTT (in min): DEXA: 228.3 ± 7.12; DEXA + HIOC: 150.0 ± 21.94, p = 0.003, Students’ t-test).

## Discussion

Our study establishes that altered BDNF–TrkB signaling in the adult ENS is an important mechanism through which the stress-induced glucocorticoid pathway manifests GI dysmotility. Our study is the first to interrogate the nature of the *Bdnf* isoforms present in the post-natal gut of juvenile, adult, and aging mice, and establishes that at all ages, the stress-responsive (hence glucocorticoid-responsive) isoforms *Bdnf IV* and *VI* together account for the majority of *Bdnf* isoforms expressed in the gut wall. Further, by demonstrating the localization of BDNF and TrkB-GFP expression in Hu-expressing myenteric cells in the adult murine gut wall, we show that the BDNF–TrkB crosstalk is neuronal in nature. Since stress increases glucocorticoid levels, and glucocorticoids downregulate the expression of *Bdnf-IV* and *Bdnf-VI* in other systems (Suri & Vaidya, 2013), we tested and found that both stress and glucocorticoid receptor (GR) agonism using a synthetic GR agonist dexamethasone significantly downregulated expression of specific *Bdnf* isoforms in the adult LM-MP. Additionally, we observed that this was associated with a significant slowdown of intestinal motility. By demonstrating that all myenteric cells ubiquitously express GR, we show that these cells are responsive to increases in glucocorticoid levels. Downstream of GR-signaling, we modulated the activity of BDNF-TrkB signaling through ENS-specific reduction of TrkB, demonstrating that this results in slowed whole gut motility, while not effecting overall enteric neuron density in the myenteric plexus. Finally, we established the involvement of BDNF-TrkB signaling in glucocorticoid-mediated GI dysmotility by showing that TrkB agonism significantly improves motility in a mouse model of dexamethasone-induced GI dysmotility. These results together show that elevated GR activation, an important down-stream effect of physiological stress, reduces the expression of stress-responsive *Bdnf* isoforms to cause GI dysmotility, an effect that can be ameliorated through stimulating TrkB signaling.

Prior studies have shown that stress and the stress hormone corticotrophin-releasing factor (CRF) in the central nervous system induce a slowdown of gastric, small intestinal, and whole gut transit (Lobo *et al*, 2023; Zhao *et al*, 2021). Through stimulation of the HPA axis, stress causes increased release of the glucocorticoid cortisol, the effect of which can be mimicked by using the GR agonist dexamethasone (Suri & Vaidya, 2013). Similar to our findings, the Thaiss lab demonstrated that both stress and dexamethasone increase whole gut transit time of orally-gavaged carmine red dye – suggesting a slowdown of GI motility (Schneider *et al*., 2023). This may appear at odds with observations of stress-induced increased defecation in thanatosis (Rogers & Simpson, 2014) and studies on stress-induced increases in colonic motility from the Tache and Vanner groups (Miampamba *et al*, 2007; Reed *et al*, 2016; Wang *et al*, 2007). However, this can be reconciled by the fact that the latter studies specifically focused on stress’s effects on colonic motility and did not consider the effects on upper or overall GI motility. Indeed, another study from the Tache group (Stengel *et al*, 2011) showed that stress slowed down gastric motility while increasing colonic motility. Since the whole gut transit assay reflects dye progression from stomach to expelled feces, we reason that the prolongation in dye transit time in our study and in the Schneider et al. study supports the conclusion that stress impairs gastric and small intestinal motility, and that this delay likely outweighs any acceleration of colonic motility which results in overall slower GI transit.

We found that both a single dose of dexamethasone and five days of restraint stress impaired gastrointestinal motility. However, while dexamethasone significantly reduced the expression of both stress-responsive *Bdnf IV* and *VI* isoforms, restraint stress selectively reduced *Bdnf IV* expression alone. This indicates that dexamethasone may exert a stronger suppressive effect on *Bdnf* expression than stress-induced elevations in endogenous cortisol. Consistent with this, dexamethasone has a substantially higher affinity for the glucocorticoid receptor than cortisol, making it a more potent GR agonist (Ballard & Ballard, 1972; Warris *et al*, 2016). It is also plausible that repeated stress led to habituation of mice, leading to a reduced effect on *Bdnf*-*VI* expression.

While our bulk RNA-sequencing analysis identified the predominant *Bdnf* isoforms, the gene’s inherently low expression at steady state likely limits the sensitivity of this approach in capturing all isoforms. This limitation is illustrated by the absence of the *Bdnf-I* isoform (whose expression is unaffected by stress) in our transcriptomic dataset, despite its detection by isoform-specific qRT-PCR assays. Notably, the absence of *Bdnf-IV*—the sole glucocorticoid-sensitive isoform downregulated by restraint stress—in aging murine LM-MP tissues may hold important implications for age-related susceptibility to stress-induced dysmotility. Investigating how these isoforms respond to restraint stress in aged mice will be the focus of future studies.

We employed both *in vivo* dexamethasone exposure and our established *ex vivo* culture system (Becker *et al*, 2013) to investigate the impact of glucocorticoid signaling on *Bdnf* isoform expression. Our findings demonstrate that the *ex vivo* model faithfully recapitulates the *in vivo* effects of dexamethasone, establishing it as a robust platform for both assessing how BDNF protein abundance is modified in response to dexamethasone exposure as well as for future studies examining how diverse exposomes influence the ENS and its surrounding cellular environment.

It is important to note that while stress-mediated changes in glucocorticoid signaling is an important mechanism through which stress manifests dysfunction, stress also drives changes in other hormones in the viscera, which include changes in adrenaline and in the peripheral release of corticotrophin releasing factor from mast cells (Kempuraj *et al*, 2017; Pace & Myers, 2023). It is currently unclear how these alternate stress-mediated pathways affect enteric BDNF–TrkB signaling, and future studies will focus on better understanding this interplay. In addition, future studies will test whether the glucocorticoid-mediated changes in the expression of *Bdnf-VI* and other isoforms occur differentially in an age-associated manner.

*Ntrk2* encodes two TrkB isoforms: a full-length TrkB.FL that faithfully carries out canonical BDNF signaling, and a truncated TrkB.T1 isoform that does not do so due to the lack of a functional intracellular kinase domain required for downstream signal transduction (Eide *et al*., 1996). As the TrkB.T1 receptor maintains the capacity to bind BDNF, competitive binding of BDNF to Truncated TrkB.T1 isoform serves to limit the availability of BDNF for the full length TrkB.FL isoform, and this quenching of available BDNF through TrkB.T1 modulates TrkB.FL-dependent intracellular signaling (Biffo *et al*., 1995). In the ENS-specific conditional TrkB mutant model mediated by excision of exon 15 (Chen *et al*., 2005; Rios-Pilier & Krimm, 2019), we observed a significant reduction in expression of transcripts encoding the TrkB.FL isoform, while abundance of transcripts encoding TrkB.T1 remained unchanged. As a result of the isoform proportion shifting even more strongly in favor of the truncated isoform, the BDNF’s downstream signaling through TrkB.FL is further reduced, leading to the observed slowing of intestinal motility.

The *Wnt1*-cre transgenic model deployed by us to drive the conditional mutation of *Ntrk2* gene is effective for the neural crest-derived neurons and glial cells of the adult enteric nervous system. We have recently shown that only half of all neurons in the adult murine small intestinal ENS are derived from the *Wnt1*-expressing neural crest (Kulkarni *et al*., 2023). This suggests that the conditional mutation of *Ntrk2* is active only in the neural crest-derived subset of ENS neurons. Thus, it is plausible that if TrkB is expressed by mesoderm-derived enteric neurons, their expression will remain unchanged. Yet, the fact that we observed a significant reduction in abundance of TrkB.FL encoding transcript and significant GI dysmotility in the *Wnt1-*cre:TrkB^fl/fl^ mice shows the relevance of TrkB expression in the neural crest-derived neurons to GI motility. Notably, exon 15 is the last shared exon of TrkB.FL and TrkB.T1 transcripts (Tomassoni-Ardori *et al*., 2019). While the specific effect of this knockout on TrkB.T1 isoform is unclear, the lack of transcriptional response and its functional role as a dominant negative inhibitor of BDNF suggest that the slowing of intestinal motility is due to loss of full-length TrkB.FL isoform in neural crest-derived neurons of the ENS.

The choice of the *Wnt1*-cre line for testing the effect of conditional Ntrk2 mutation on GI function was also necessitated by the fact that there is currently no available transgenic cre line specific to a large proportion of ENS cells. Other transgenic lines that have been used by other investigators include the *Hand1*-cre mouse line (Schneider *et al*., 2023) that can label non-ENS cells in the gut wall, or the Baf53b-cre line (Morarach *et al*, 2021) that will also cause loss of TrkB isoforms in central nervous system (Zhan *et al*, 2015), thus making it difficult to segregate the central nervous system’s effects from the local ENS effects. Future studies would require us to identify the nature of the ENS neurons expressing *Bdnf* and *Ntrk2* and utilize inducible cre systems to interrogate how loss of these genes by specific neuronal subpopulations in the ENS affect GI function.

Alterations and mutations in *Bdnf* and *Ntrk2* have been associated with GI dysmotility (Bonfiglio *et al*, 2021; Linsalata *et al*, 2023; Weaver *et al*, 2022), and exogenous supplementation of BDNF has been observed to have a prokinetic effect in patients (Chen *et al*., 2014; Coulie *et al*., 2000). However, the cellular and molecular mechanisms that regulate BDNF levels in these diseases of gut-brain interactions (DGBI) that are often associated with aberrant stress responses were unknown. Our study, which identifies the neuronal nature of BDNF and TrkB, characterizes the major *Bdnf* isoforms in the LM-MP tissue at three different ages, and studies the effect of stress and GR signaling on *Bdnf* isoform abundance, is the first to establish the mechanism through which this clinically relevant biomolecule is altered in the gut. Previously, it has been shown that BDNF amplifies the responsiveness of enteric neurons to stimuli (Boesmans *et al*., 2008), suggesting that this neurotrophin regulates the ‘*gain*’ of TrkB-expressing neurons. Here, using transgenic TrkB-GFP mice, we have estimated this population to be ∼16% of all adult small intestinal Hu^+^ myenteric cells. With our data showing that increasing TrkB signaling in dexamethasone-treated mice improves GI motility, this suggests that stress-mediated downregulation of BDNF reduces the sensitivity or responsiveness of TrkB-expressing myenteric neurons, leading to dysmotility. Together, this identifies that TrkB is an important therapeutic target for the amelioration of GI dysmotility in DGBI patients.

Thus, by studying an important mechanism through which stress-associated changes in glucocorticoid signaling alters the BDNF–TrkB crosstalk in the adult ENS, our study identifies an important pathway through which stress causes GI dysmotility and provides a putative therapeutic target which can be used for improving GI motility in many DGBI patients.

### Transcriptomic Data and Code availability

All the transcriptomic data has been deposited in GEO GSE284108 and the code for analysis is available at https://github.com/jaredslosberg/timecourse_lmmp_bulkrnaseq/

## Acknowledgements

We acknowledge Dr. Xia Qian (Dept of Pathology, BIDMC) and Mrs. Vilmonse (Eva) Csizmadia (Dept of Anesthesia, BIDMC) for their help. This work was supported by funding from NIA R01AG066768, R21AG072107, Diacomp Foundation (Pilot award Augusta University) and Pilot grant from the Harvard Digestive Disease Core (SK). This work was also supported by funding from the Maryland Genetics, Epidemiology, and Medicine training program sponsored by the Burroughs Welcome Fund (JS), NIGMS grant T32GM148383 (JS), and from Walter Benjamin Fellowship (528835020) from Deutsche Forschungsgemeinschaft (PS).

Extended Figure 2 – 1: Gene and isoform expression of *Bdnf* by age. (A) *Bdnf* expression is detected in 1-month, 6-month, and 17-month-old mice. Isoform-level abundance estimates were collapsed to gene-level after inter-library normalization with edgeR (TMM method). Across all ages, average expression of *Bdnf* is 0.38 ± 0.02 TPM. Independently, gene-level differential expression across age was conducted with DESeq2 (counts ∼ age + RIN + batch, Wald test) and *Bdnf* was not identified as differential by age across any comparison. (B) Abundance estimates of individual *Bdnf* isoforms identify the e6 isoform (ENSMUST00000111045) as the majority isoform in each condition. The average expression of this isoform is 0.32 ± 0.08 TPM. Other lowly detected isoforms include e5 (ENSMUST00000176893), e4 (ENSMUST00000111050), e3 (ENSMUST00000111046), and e2 (ENSMUST00000111047), while remaining isoforms were not detected. Differential isoform usage by age was tested via the DEXSeq wrapper within IsoformSwitchAnalyzeR (Likelihood ratio test, full model: ∼ sample + exon + age:exon, reduced model: ∼ sample + exon). No *Bdnf* isoforms were identified as having significant differential usage by age. (C) Isoform level expression was transformed to isoform fraction (IF) measures by age (for each age, sum of IF’s is 1).

