## Supplementary figures and images for "Reduced enteric BDNF-TrkB signaling drives stress-dependent glucocorticoid-mediated GI dysmotility"

### Suppl. Fig 1

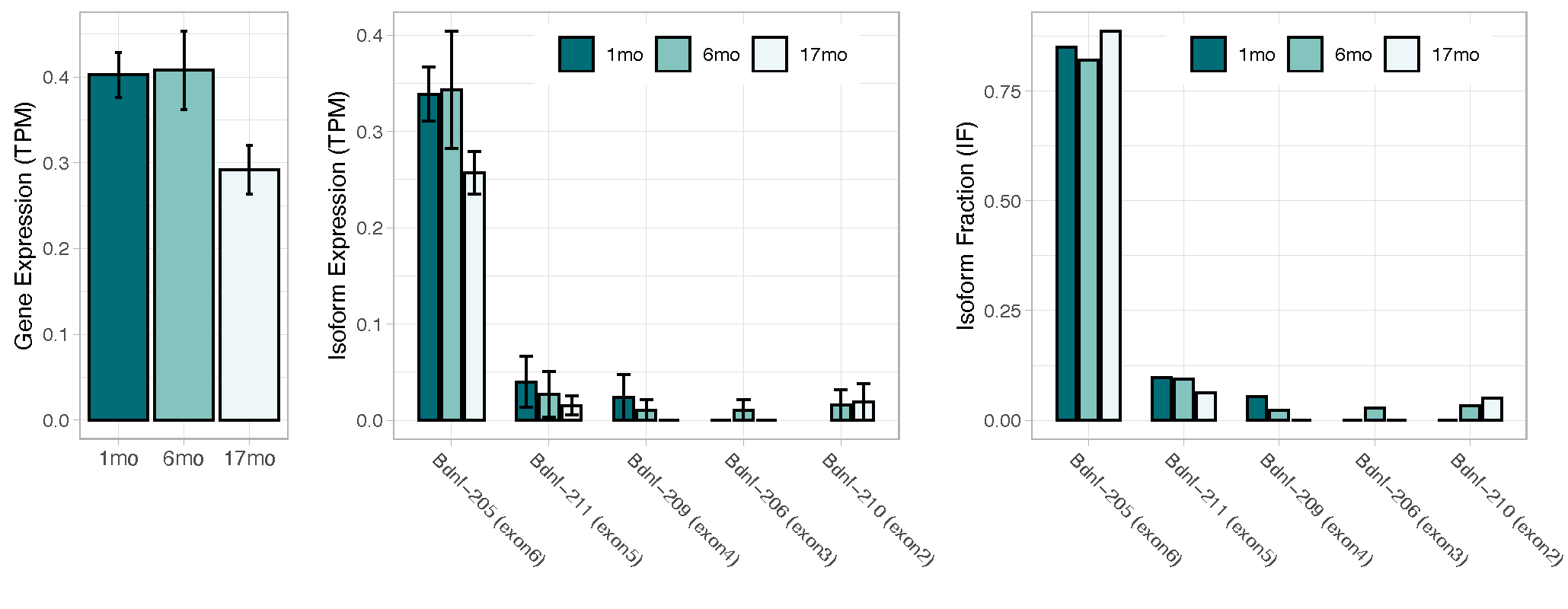
